# A rolling circle-replicating plasmid as an Inovirus phage satellite

**DOI:** 10.1101/2023.11.28.569023

**Authors:** Nicole E Schmid, David Brandt, Claudia Walasek, Clara Rolland, Johannes Wittmann, Mathias Müsken, Jörn Kalinowski, Kai M Thormann

**Author notes:** corresponding author: Kai M. Thormann, Institute for Microbiology and Molecular Biology, Justus-Liebig-Universität Gießen, Heinrich-Buff-Ring 26, D-35393 Gießen, Germany.

## Abstract

Bacterial viruses (phages) are potent agents of lateral gene transfer and thus are important drivers of evolution. A group of mobile genetic elements (MGEs), referred to as phage satellites, exploit phages to disseminate their own genetic material. Here we isolated a novel member of the genus *Inovirus*, *Shewanella* phage Dolos, along with an autonomous rolling circle-replicating plasmid, pDolos. Dolos causes a chronic infection in its host *Shewanella oneidensis* by phage production with only minor effects on the host cell proliferation. When present, plasmid pDolos hijacks Dolos functions to be predominantly packaged into phage virions and released into the environment. pDolos can disseminate further genetic material encoding, e.g., resistances, fluorophores, and metabolically active proteins, to host cells sensitive to Dolos infection. Given the rather simple requirements of a plasmid for takeover of an inovirus, the wide distribution of phages of this group and the broad spectrum of rolling circle-replicating plasmids, we speculate that similar phage-satellite systems are common among bacteria.

## Introduction

Lateral gene transfer between species is a main driver of evolution. A highly diverse range of mobile genetic elements (MGEs) has been identified to translocate DNA, comprising agents such as integrative and mobilisable elements (IMEs), integrons, transposons, insertion sequence (IS) elements, integrative and conjugative elements (ICEs), plasmids and viruses ^1,2^. Viruses of bacteria (referred to as bacteriophages or phages) are effective shuttles of genetic material and are themselves exploited by satellite mobile elements ^3^. This group of mobile elements includes phage satellites (such as the well-studied P4/P2 system), phage-inducible chromosomal islands (PICIs) and phage-inducible chromosomal island-like elements (PLEs) ^4–8^. All of these elements reside within the host genome and are unable to produce infectious virions on their own, but instead rely on a so-called helper phage for lateral dissemination ^9–11^. Compared to the hijacked phage, these phage-exploiting elements have substantially smaller genomes harbouring phage-like genes for replication, propagation and shuttle-phage takeover. They may also encode virulence factors or systems to limit further phage predation ^8,9,12^, which can provide an advantage for their bacterial host. Phage-hijacking elements are highly abundant in a wide array of bacterial species, however, their diversity and properties so far remain largely unexplored ^10,11,13^.

Phages exhibit a huge diversity in morphology and genomes. Among them, the family of *Inoviridae* consists of several subtypes, one of which is the genus *Inovirus*, comprising well-known representatives such as the *Escherichia coli* Ff phages (M13, fd and f1), *Pseudomonas aeruginosa* phages Pf1 and Pf3, or *Vibrio cholerae* phage CTX ^14,15^. These phages are characterized by a filamentous morphology, a circular single-stranded DNA genome and an intricate life cycle. Inoviruses can be integrated into the host chromosome but may also remain freely in the cytoplasm. Commonly, phages of this group do not lyse and kill their host but cause a chronic infection. Upon entry into the host cell the circular single-stranded phage genome will be converted by host enzymes into double-stranded circular DNA (dsDNA). This dsDNA phage chromosome undergoes rolling-circle replication, and the single-stranded DNA, stabilized by single stranded DNA-binding (SSB) proteins, is recognized trough binding of a phage protein to a specific recognition DNA structure. This complex is fed into the intricate virus-encoded extrusion machinery, which releases the SSB units and covers the DNA by phage coat proteins during transport (see ^14,15^). This way, the host cells can produce and release large amounts of phage particles without lysis, which, however, often causes reduced growth due to loss of cellular resources required for the production of phage building blocks and DNA. Inoviruses have been shown to affect their host in various ways, e.g. by altering pathogenicity and biofilm formation ^14–20^. Despite numerous diverse inoviruses present among bacteria and archaea (Roux et al, 2019), only some have been studied in more detail.

Here, we describe the identification of Inovirus Dolos, which infects the gammaproteobacterium *Shewanella oneidensis* MR-1 and some related species. Along with the phage, we isolated an autonomously replicating plasmid, which hijacks *Shewanella* phage Dolos as a phage satellite for lateral dissemination.

## Results

### Isolation and characterization of Inovirus Shewanella Phage Dolos

In a study aimed at the isolation and identification of phages infecting *Shewanella oneidensis* MR-1 ^21^, water and sediment samples of limnic environments were sampled. One of the enrichments from a fresh water sediment yielded a phage isolate characterized by small turbid plaques (**Fig. 1D**). Electron microscopy on phage particles, which were enriched three times via single plaques and purified by density gradient centrifugation, identified a filamentous phage of about 1 µm in length and 7.5 nm in diameter (**Fig. 1C**). The distinct morphology in concert with phage genome sequencing identified the isolate as a novel representative of the *Inoviridae* group, which was named *Shewanella* phage Dolos (henceforth referred to as Dolos, **Supplementary Fig. S1**). Dolos possesses a single-stranded DNA (ssDNA) genome of 8146 nt in length containing 11 predicted putative open reading frames (ORFs) (Genbank accession: OP867012.1). According to homologies and protein domain analysis, potential functions for genome replication, phage assembly and phage extrusion were predicted for 7 of the gene products (ORF1-ORF7). The ORFs 8, 10 and 11 encode proteins of unknown function, Orf9 has some homologies to an antitoxin of a Maz-like toxin-antitoxin system (**Fig. 1A** and **Supplementary Table S1**). Sequencing of phage-infected *S. oneidensis* cultures showed that no Dolos integration into the host chromosome occurs.

**Figure 1.**
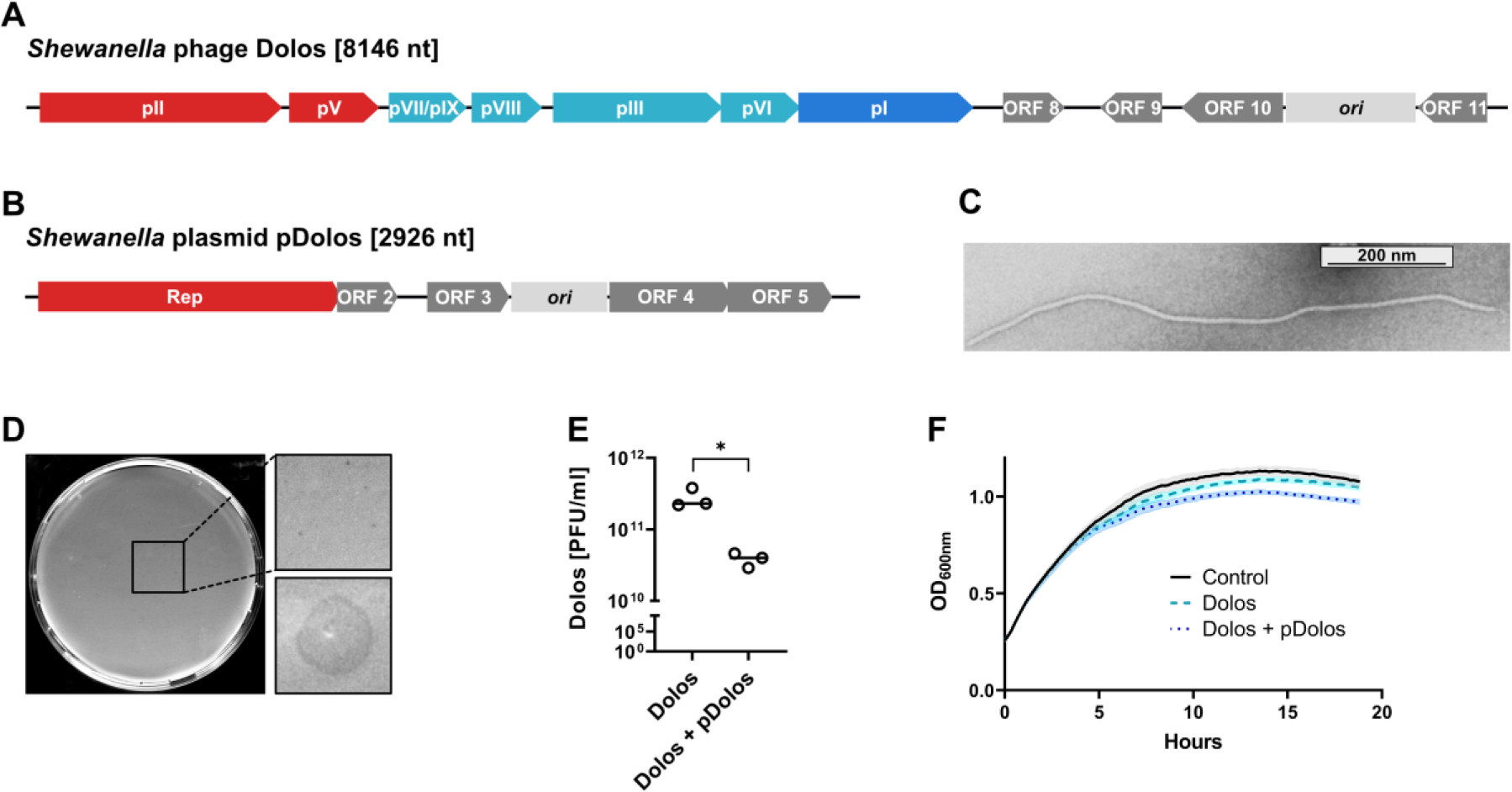
Characterization of *Shewanella* phage Dolos and the co-isolated rolling circle plasmid pDolos. **A, B.** Predicted genetic maps of *Shewanella* phage Dolos and the plasmid pDolos. The putatively identified gene products are indicated. See also **Supplementary Table S1**. Red, proteins functioning in replication; turquoise, structural proteins; blue, secretion protein; dark grey, proteins of unknown or putative function; light grey, putative phage/plasmid origin regions. **C.** Electron microscopic image of filamentous phage Dolos. Phage containing supernatant was purified by cesium density gradient centrifugation and negatively stained for imaging. **D.** Plaque assays demonstrating turbid plaque morphology of inovirus Dolos. **E.** Phage concentration of the phage-containing supernatants of the host strain infected with Dolos or both Dolos and pDolos. Cells were infected with Dolos or Dolos/pDolos at mid-exponential phase at a MOI of 0.1. 24h after infection the supernatant of infected cells was sterile filtered and the PFU/ml calculated. The asterisk represents p < 0.05 (t-test). **F.** Growth of *S. oneidensis* cultures infected with phage containing supernatant Dolos or Dolos/pDolos. Cells were infected during mid-exponential phase at a MOI of 0.1 and the OD_600_ was measured subsequently for 18h. Shown are the results of three independent experiments.

Under the conditions tested, phage extrusion started about 120 min after infection. As typical for inoviruses, infection of *S. oneidensis* MR-1 with Dolos at an MOI (multiplicity of infection) of 0.1 only had a minor negative effect on growth of the culture despite the production of up to 10^11^ phage particles occurring in the supernatant of stationary cultures (**Fig. 1E, F**). The phage particles remained stable between pH 4 and pH 12 and were inactivated by extended exposure to temperatures higher than 45 °C (**Supplementary Figure S2**). Among 27 different isolates and species of *Shewanella* (as well as *E. coli* MG1655, *Pseudomonas putida* KT2440 and *Vibrio cholerae* El Tor N16961) Dolos infected, in addition to *S. oneidensis* MR-1, two *S. baltica* isolates, and thus appears to have a rather narrow host range (**Supplementary Table S2**). The presence of the two active prophages in *S. oneidensis* MR-1, LambdaSo and MuSo2 did not affect the ability of Dolos to infect the cells. However, to avoid any interference caused by cell lysis and /or plaque formation, we continued our studies using a strain lacking both active prophages (MR-1 ΔLambdaSoΔMuSo2). In the following, this strain will be referred to as wild-type *S. oneidensis* for simplicity.

### A rolling circle plasmid (pDolos) co-isolated with Dolos

Isolation of virion-integrated DNA from phage-containing supernatants and subsequent sequencing showed the presence of a second genetic element in addition to the chromosome of phage Dolos. This element was identified as a plasmid of 2926 nt, which was also not found to be inserted into the host chromosome (Genbank accession: OR865871). The plasmid, which we termed pDolos, is predicted to encode five proteins (**Fig. 1B**). One of the encoded gene products (ORF1, 359 aa), belongs to the Rep proteins, which are responsible for replication of rolling-circle plasmids ^22^. The other gene products are of unknown function (ORF2, 47 aa), a putative transcriptional repressor (Orf3, 81 aa), a putative antitoxin of a type II class toxin-antitoxin system (ORF4, 130 aa), and a putative ssDNA-binding protein (ORF5, 109 aa). None of these proteins had any apparent homologies to known phage genes.

In the following, we checked whether stability of pDolos depends on presence of phage Dolos - or *vice versa*. Phage Dolos without the plasmid could be readily obtained by screening single plaques after dilution of Dolos-containing culture supernatants. The phage without the plasmid did not exhibit any negative effect on phage propagation and plaque formation, indicating that Dolos does not require any functions of the plasmid. To determine the stability of pDolos without the phage, a kanamycin-resistance cassette was integrated into the plasmid behind ORF5. The resultant construct was electroporated into *S. oneidensis*, yielding kanamycin-resistant colonies. Plasmid-containing strains were then cultivated three consecutive times to stationary growth phase without the presence of kanamycin. Subsequent quantification of colony-forming units showed that pDolos was stably maintained also in the absence of selective pressure or presence of the phage (**Fig. 2D**). We therefore concluded that pDolos, the second genetic element isolated from the phage-containing supernatant, is a plasmid autonomously replicating via rolling-circle mechanism. However, no free infectious particles nor plaque formation was observed when pDolos was present without the phage, indicating that the plasmid transfer between cells relies on functions provided by phage Dolos.

**Figure 2:**
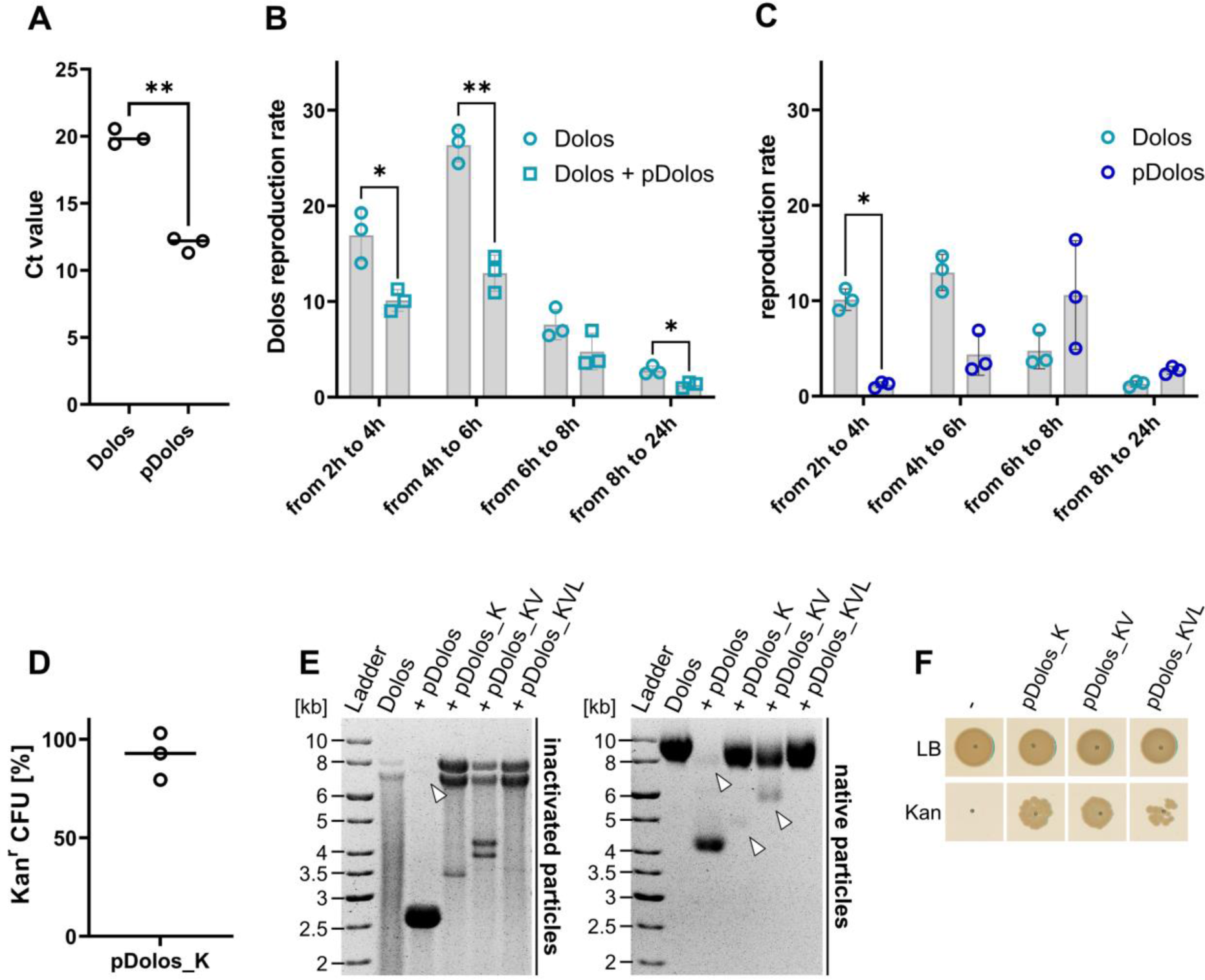
pDolos acts as phage satellite of phage Dolos. **A.** qPCR-derived Ct values of Dolos genome and pDolos genome in an outgrown supernatant containing phage Dolos and satellite pDolos. The difference corresponds to a plasmid/phage ratio of about 250. **B and C.** Calculated reproduction rate of a time-serial infection at a MOI of 0.1. Samples for qPCR analysis were taken 2h, 4h, 6h, 8h and 24h after infection of mid-exponential cells with only phage Dolos or Dolos together with satellite pDolos. *, p < 0.05; **, p < 0.01 (t-test). **B.** The reproduction rate of Dolos over time was estimated using phage containing supernatant Dolos and the mixed infectious supernatant containing Dolos and the satellite pDolos. **C.** The reproduction rate of satellite pDolos compared to that of phage Dolos**. D.** Percentage of kanamycin-resistant cells harboring plasmid pDolos containing the appropriate resistance gene (pDolos_K). Cells harboring the plasmid were cultivated without kanamycin pressure over three days. **E.** Agarose gel showing DNA after separation of heat-inactivated particles (free DNA; left panel) and native particles (right panel) by electrophoresis. To visualize DNA of native particles, staining was performed after disruption of particles by NaOH. As indicated the samples contained solely phage Dolos or, in addition, pDolos constructs of increasing sizes (pDolos_K; pDolos_KV and pDolos_KVL). The DNA often appeared as a double band on the gel. **F.** Transduction of pDolos variants by phage Dolos. Cells were infected with an outgrown supernatant prepared from cells containing different pDolos constructs, which were then infected by Dolos. 24h after transduction with the prepared infectious supernatant, cells (OD_600_ 1) were spotted on LB_Km_ plates. Colony growth was imaged after 24h of incubation.

### pDolos acts as a phage satellite

The co-isolation of Dolos and pDolos from phage-containing supernatants strongly suggested that pDolos is packaged and released from the cell, either as an independent particle or together with the phage genome in a single virion. Due to the extrusion mechanism of inoviruses ^15^, we hypothesized that packaging of the significantly smaller plasmid DNA leads to formation of an accordingly smaller particle. We therefore separated the virions of a phage-containing supernatant by agarose gel electrophoreses followed by disruption of the virions’ protein shell to release the DNA. We found that in the presence of pDolos, a distinct secondary DNA band was present at a position corresponding to notably smaller virion size in addition to the larger Dolos particle (**Fig. 2E**). PCR analysis of the DNA revealed that the released particles exclusively contained either Dolos or pDolos DNA. The results strongly indicated that pDolos is packaged and released by the Dolos machinery and thus acts as a phage satellite. Notably, phage and plasmid DNA released before gel separation (Fig. 2E, left panel) appeared as a double band and may represent different structures of the single-stranded DNA.

In addition, we determined the reproduction rate of Dolos and pDolos within a culture by qPCR. Notably, additional production of the pDolos virion had a rather minor effect on growth of the *S. oneidensis* host cell culture (**Fig. 1F**). During early growth (from 2 to 4 and from 4 to 6 hours), reproduction of phage Dolos was significantly higher than that of the satellite plasmid pDolos (**Fig. 2C**), but presence of pDolos already significantly decreased the reproduction of Dolos (**Fig. 2B**). At later stages (6 to 8 and 8 to 24 hours), the reproduction of pDolos increased and became higher than that of the phage. In the supernatant of an outgrown culture containing both phage and its satellite, the amount of the latter exceeded that of the phage itself by a factor of about 250. Accordingly, the number of Dolos phage particles in the supernatant of an outgrown colony decreased by one order of magnitude (**Fig. 1E; 2A**).

### Diverse genes can be transduced by Dolos/pDolos

The results indicated that pDolos effectively hijacks the packaging and extrusion apparatus of Dolos. We next asked if the virions with the plasmid are infective and if further genes can be delivered to cells. To answer this, we used the already present version of pDolos that carries an additional kanamycin resistance cassette. In addition, versions of pDolos were constructed that harboured the gene encoding the yellow fluorescent protein mVenus and the *lacZ*’ gene from *E. coli*, thereby also considerably increasing the size of the plasmid (pDolos_K, 4,015 bp; pDolos KV, 5,533 bp; pDolos_KVL, 9,850 bp). Electrophoretic separation of the formed particles showed that also the larger variants of pDolos are successfully packaged into virions, which accordingly increased in size (**Fig. 2E**). Subsequent infection experiments resulted in plasmid-bearing cells that were resistant to kanamycin (all variants) and displayed yellow fluorescence (in pDolos_Km^r^_mVenus and pDolos_Km^r^_mVenus_LacZ’). Plasmid and kanamycin resistance transfer also occurred to other *Shewanella* species (**Supplementary Fig. S3**). These results show that pDolos can carry additional genes that are subsequently transduced into host cells of bacterial species sensitive to Dolos infection.

## Discussion

Phages are important drivers of lateral gene transfer ^23^ and are themselves exploited by genetic elements, so-called phage satellites, for their dissemination. A plethora of diverse phage satellites has been identified throughout different species of bacteria ^13^ to which we now add a very simple phage hijacking system that uses an inovirus for intercellular transfer. This phage satellite, pDolos, is an autonomously rolling circle-replicating plasmid, which, notably, is not integrated into the host chromosome, in contrast to all other phage satellites characterized so far. Hijacking an Inovirus is a very efficient way of plasmid dissemination as the host cell is not lysed during this process but instead is turned into a producer of phages and phage satellites with comparably little effect on host proliferation itself.

The simplicity of the inovirus packaging system has been recognized early on, and vectors based on the *E. coli* Ff inoviruses, mainly M13, were constructed that allowed the packaging of single-stranded DNA ^24–26^. To this end, the DNA containing phage origin and packaging signal was inserted into regular cloning vectors, so-called phagemids, which was sufficient for production of ssDNA-containing virions by exploiting the functions of a M13 helper phage. It is thought that phage satellites may have evolved by different mechanisms, *de novo* evolution, gene acquisition and (gene) reduction of temperate phages ^8^. It cannot be excluded that pDolos originates from an Inovirus that has drastically reduced its genome until no recognizable phage proteins remained. However, in the system found in this study a gain-of-function of a rolling-circle plasmid that allows phage hijacking is easily conceivable as the sole requirement lacking is a short gene region (60 bp in *E. coli* phage f1), which serves as the packaging signal ^15,27–29^. As Inoviruses, rolling circle-replicating plasmids are abundant in nature and can be found in Gram-positives, Gram-negatives and Archaea ^29^. We therefore speculate that plasmid dissemination by inoviruses may be rather common. However, if other already identified rolling-circle plasmids are similarly able to be transduced by inoviruses remains to be seen.

The function of most of the gene products encoded on the pDolos remains elusive so far. One of the predicted gene products is vaguely reminiscent to an antitoxin (NdoAI/MazE) of an mRNA-cleaving type II toxin-antitoxin (TA) system ^30,31^. Generally, TA systems have been implicated in mediating plasmid stability but also in persister cell formation and phage defence (reviewed in ^32–34^). pDolos thus may be beneficial in host cell proliferation and survival, which would equally benefit its own stability and spreading. We observed a strong decrease in Dolos production in the presence of its phage satellite pDolos, which may rather simply be a matter of competition for ressources, e. g., the proteins required for packaging. Notably, the timing of maximal offspring production appears to be different for phage Dolos and its satellite. Synthesis of Dolos DNA peaks during at early time points during culturing and then drops, as also observed earlier for *E. coli* phage f1 ^35^. In contrast, synthesis of pDolos DNA is highest at later time points, suggesting an order of production with Dolos being released first and pDolos produced later. It can be hypothesized that this may benefit dissemination of the plasmid as more cells are likely already harboring phage Dolos before advent of its satellite. The additional proteins encoded on pDolos may also be required for a more efficient takeover of its shuttle phage. Phage-satellite interaction and the potential effect of the plasmid-encoded gene products on host or phage proliferation is currently under investigation.

## Materials and Methods

### Bacterial growth conditions and measurement of planktonic growth

*E. coli* strains were cultivated at 37 °C, all other strains were cultivated at 30°C. Strains were grown in or on LB medium with the exception of some *Shewanella* isolates (as indicated) that were cultivated in or on 4M Medium ^36^ supplemented with 0,5 % (w/v) casamino acids. For plates, the appropriate media were supplemented with 1.5 % (w/v) agar. If necessary, media were supplemented with kanamycin at a concentration of 50 mg·ml^-1^. Bacterial planktonic growth was monitored by measuring the OD_600_ of a 200-µl culture in 10-minute intervals in a microtiter plate using a plate reader (Tecan INFINITE M NANO+; Tecan; Männedorf; Switzerland).

### Virion isolation from limnic samples

Virions were isolated like described earlier ^21^. The mixed Dolos- and pDolos-containing supernatant was obtained after the first step of virion isolation by culturing *S. oneidensis* MR-1 ΔLambdaSo ΔMuSo2 with a limnic sample for 24 h. Pure Dolos phage-containing supernatant was reached after two consecutive singling-out incubations of single-plaques with a liquid culture of *S. oneidensis* MR-1 ΔLambdaSo ΔMuSo2.

### Genome sequencing, assembly and annotation

Extracted phage DNA was sequenced using Illumina (San Diego, USA) and Oxford Nanopore (Oxford, UK) technology. For Oxford Nanopore sequencing, the Rapid Barcoding kit (SQK-RBK004) and a R9.4.1 flowcell were used. Sequencing was done on a GridION machine. Illumina libraries were prepared using the NanoLT workflow and sequenced on a MiSeq machine. After sequencing, Unicycler v0.4.6 ^37^ was used for hybrid genome assembly and the assembled contigs were annotated using prokka v1.14.5 ^38^ with the supplementation of HMMs from the PHROG database and refined using AlphaFold structure predictions ^39,40^. Unless specified otherwise, tools were used with standard settings. Annotated genome sequences were deposited at Genbank.

### Construction of pDolos plasmid variants

Bacterial strains, plasmid and primers used in this study are listed in **Supplementary Tables 3 and 4**. For the DNA manipulation of pDolos, the plasmid was linearized via PCR by generating forward and reverse primers that were adjacent to each other at the same position in the plasmid. Using these primers, the whole plasmid was amplified by using the Thermo Scientific Phusion High-Fidelity DNA-Polymerase (Thermo Fisher Scientific Inc.). Constructs of pDolos were generated with the Gibson assembly method ^41^. The resultant constructs were gel purified and then transformed into *S. oneidensis* MR-1 strains by electroporation.

### pDolos stability assay

Stability of plasmid pDolos was determined over three consecutive re-growth phases in LB medium of cells containing plasmid pDolos_K. In mid-exponential growth during the third re-growth without kanamycin pressure, cells were diluted and plated on LB agar plates with or without kanamycin. Colony forming units (CFU) were determined after 24 h of growth. The experiment was carried out in three biological replicates.

### Transduction of pDolos constructs into Shewanella cells

To transduce virions of different pDolos variants into bacteria, mixed virion-containing supernatants were generated by infecting appropriate mid-exponential *S. oneidensis* cell cultures containing different pDolos plasmid variants with phage Dolos (MOI 0.1). 24h after infection, the virion-containing supernatants were isolated as previously described ^21^. These outgrown virion-containing supernatants then were used to transduce the different pDolos variants into *Shewanella*. To this end, mid-exponential cells were infected with these supernatants at MOI 2. 24 h after transduction, re-grown cells (OD_600_ 1) were spotted on kanamycin plates. Colony growth was imaged after 24 or 48 h of incubation depending on the used medium for culturing.

### Virion enrichment and DNA isolation

Virion enrichment and DNA isolation was conducted essentially by a variation of an established protocol (Phage DNA extraction - traditional. Texas A&M University, College Station, TX 77843 Rev. 9/21/2018). For virion enrichment, 10 ml of virion-containing supernatant was incubated overnight at 4°C with 5 ml of 30% (w/v) PEG 8000 in 3M NaCl. Particle precipitation was conducted by centrifugation of samples at 10,000 x g for 90 min. Sediments were re-suspended in 5 mM MgSO_4_ for DNA extraction or in medium for simple virion enrichment. For DNA extraction of particles, free DNA and RNA in the samples were first digested with DNAse I and RNase A (final concentration: 0.1 mg/ml) for 1 hour at 37 °C. The subsequent DNAse inactivation was carried out with a final concentration of 20 mM EDTA. For subsequent particle digestion, samples were incubated for 1h at 60°C at a final concentration of 20 µg Proteinase K and 10 % (w/v) SDS. Afterwards, virion DNA was isolated by phenol-chloroform extraction. To this end, an equal volume of phenol:chloroform (1:1) was added and the samples were inverted several times. DNA-containing water phase was separated after centrifugation at 6000 x g for 5 min and the phenol-chloroform step was repeated with the separated water phase. Following, an equal volume of chloroform was added to the samples, which were then gently mixed by inverting, and the water phases were separated after centrifugation by pipetting. DNA precipitation was conducted by adding 1/10 volume of 3M sodium acetate (pH 7.5) and 2.5 volumes of 96 % (v/v) ice cold ethanol. After storage overnight at -20°C, the DNA was precipitated by centrifugation at 10,000 x g for 30 min. After centrifugation the supernatant was removed and samples were washed twice by filling the tubes halfway with 70% (v/v) ethanol following a centrifugation at 10,000 x g for 3 min. Afterwards, tubes were opened to let the last amount of ethanol disperse and DNA was dissolved in 50 µl of water and stored at 4°C.

### Chromosomal DNA isolation of infected S. oneidensis MR-1 cells

To determine if Dolos and pDolos can integrate into the host chromosome, the chromosomal DNA of *S. oneidensis* MR-1 cells infected with a mixed Dolos- and pDolos-containing supernatant was isolated using E.Z.N.A Bacterial DNA Kit (Omega Bio-Tek). Hence, mid-exponential cells were infected with an MOI of 0.1 and 24 h after infection chromosomal DNA of 1 ml cells was isolated and sequenced.

### Quantitative PCR (qPCR)

To determine the amount of pDolos compared to the amount of Dolos, qPCR was performed. Virion DNA of Dolos-containing supernatants and of Dolos- and pDolos-containing supernatants were isolated in biological triplicates. Additionally, chromosomal DNA of cells at different stages of infection was isolated. To this end, *S. oneidensis* MR-1 ΔLambdaSo ΔMuSo2 cells were infected during the mid-exponential growth phase with an MOI of 0.1 with Dolos or with a supernatant containing mixed Dolos and pDolos virions. DNA was isolated from biological triplicates at 2 h, 4 h, 6 h, 8 h and 24 h post-infection by following the protocol of the E.Z.N.A Bacterial DNA Kit (Omega Bio-Tek) up to the column step and then performing phenol-chloroform DNA extraction (see above) on the samples. qPCR was performed by using the PowerUp™ SYBR™ Green Master Mix (Thermo Fisher Scientific Inc.), the qPCR cycler QuantStudio3 (QuantStudio^TM^) as well as the software “Design & Analysis” (QuantStudio^TM^). Ct-ratios were calculated with the formular: 2^-ΔCt^ ^42^. Infection rate was calculated by normalisation of these ratios.

### Native agarose gel electrophoresis assay

The native gel electrophoresis method developed by Bennett et al., 2011 ^43^ was used to determine differences in virion size depending on the packaged genome. For this purpose, the free DNA of outgrown virion-containing supernatant was digested by incubating the supernatant with 0.1 mg/ml DNAse I for 15 minutes at 37 °C. To separate virion DNA, 10 µl of virion-containing supernatant was incubated with 2 µl DNA loading dye (30% glycerol (v/v), 0.25% bromophenol blue dye (w/v), and 0.25% xylene cyanol FF dye (w/v) in 1x TBE) and a final concentration of 1% (w/v) SDS and heat inactivated at 80 °C for 15 min. To separate virions, 2 µl DNA loading dye was added to the 10 µl of supernatant. Samples were loaded on a 0.6% (w/v) TBE-agarose gel. Virions and DNA were separated by electrophoresis at 2.5 V / cm for 14,5 h. Visualization of free DNA was conducted by a 15 min incubation of the gel with ethidium bromide (1µg/ml). To visualize packed DNA, particles were digested using a 0.2 M NaOH bath for 45 min. Afterwards the gel was neutralized using a 0.45 M Tris HCl (pH 7.2) bath for 15 min. A second ethidium bromide staining and UV-light analysis revealed the DNA of the virions running through the gel.

### Density centrifugation, staining, electron microscopy

For preparation of the Caesium Chloride (CsCl) gradient, 300 ml of Dolos phage lysate (∼10^11^ PFU/mL) were centrifuged for 2 h at 16,000 rpm (Sorvall RC6Plus, F21S 8 × 50Y) in order to concentrate the phages. The phage was resuspended in 4 ml of LB media and purified by ultracentrifugation in a discontinuous CsCl gradient (0.5 mL CsCl solution with densities of 1.6, 1.5, 1.4, and 1.3) for 2 h at 4°C and 35,000 rpm in an SW 60 Ti rotor. Phage bands were then removed and dialyzed against SM buffer (100 mM NaCl, 8 mM MgSO_4_, and 50 mM Tris-HCL (pH 7.5) ^44^.

For transmission electron microscopy (TEM) analysis, phages were prepared as previously described^45^. Briefly, phages were allowed to adsorb onto thin carbon support films. Afterwards, they were negatively stained with 2% (w/v) aqueous uranyl acetate, pH 5.0. Samples were examined in a Zeiss EM 910 or Zeiss Libra120 Plus transmission electron microscope (Carl Zeiss, Oberkochen, Germany) at an acceleration voltage of 80 kV/120 kV at calibrated magnifications using a crossed line grating replica. Size determination was performed using ITEM Software (Olympus Soft Imaging Solutions, Münster).

### Phage spot and plaque assay

To analyse infectivity of Dolos, plaque and spot assays were conducted. To this end, 400 µl of stationary cell culture were mixed with 7 ml soft agar [0.5% (w/v)], mixed by vortexing and poured over an agar plate (agar overlay method). In case of a plaque assay, 100 µl of diluted phage-containing supernatant was mixed with the soft agar before pouring. For a spot assay, phage-containing supernatants were spotted on top of a hardened overlay plate. Plaque and spot formation was investigated after two days of incubation.

### Temperature and pH sensitivity assay

To investigate pH stress resistance of phage Dolos, 10 µl of a phage-containing supernatant (10^9^ PFU per ml) were incubated with 90 µl of LB medium adjusted to different pH values at RT. Phages were incubated in media with different pH for 24 h at RT. To test thermal stability of Dolos, virions phage-containing supernatant (10^9^ PFU/ml) was incubated for 24h at indicated different temperatures using a thermal-cycler (Eppendorf). Afterwards, supernatants were serially diluted using LB medium and 1.5 µl of each dilution was spotted on top of agar overlay plates. Plaque formation was investigated after two days of incubation. pH and temperature sensitivity were determined in three independent biological replicates.

### Fluorescence microscopy

*S. oneidensis* MR-1 ΔLambdaSo ΔMuSo2 cells transduced with different pDolos constructs were analysed via fluorescence microscopy to examine fluorophore presence in transduced cells. To this end, stationary cells were diluted to an OD_600_ of 0.02 and incubated until mid-exponential growth phase. 2 µl of cell culture were spotted on an agarose pad [1% (w/v) agarose in LM medium (10 mM HEPES, pH 7.5; 100 mM NaCl; 0.02% yeast extract; 0.01% peptone; 15 mM lactate)] and images were recorded using a Leica DMI 6000 B inverse microscope (Leica, Wetzlar, Germany) fitted with an sCMOS camera and a HCX PL APO 100x/1.4 objective. VisiView software (Visitron Systems, Puchheim, Germany) was used for image aquisition. Image processing was carried out via the Fiji tool ^46^.

### Graph design and statistics

Graphs were designed using GraphPad PRISM (Graphpad Software, Inc). Depending on the data, multiple paired and unpaired t-test were also performed with the PRISM software (*, p < 0.05; **, p < 0.01).

## Acknowledgements

The authors are grateful to Susanne Brenzinger and Ana Rita Brochado from the University of Würzburg for testing Dolos infection on *Vibrio*. We also thank Ina Schleicher for technical support in EM sample preparation and acknowlegde the NGS Core Facility at Bielefeld University for providing the sequencing data. Finally, we are grateful to Callypso Pellegri and Laetitia Houot from the University of Aix-Marseille for their helpful hint on the native gel electrophoresis method and to Eduardo Rocha for the insightful comments on the manuscript.

## Conflict of interest

The authors declare no conflict of interest.

## Notes

### Competing Interest Statement

The authors have declared no competing interest.

## References

1. Frost, L. S., Leplae, R., Summers, A. O. & Toussaint, A. Mobile genetic elements: the agents of open source evolution. Nat Rev Microbiol 3, 722–732 (2005).

2. Weisberg, A. J. & Chang, J. H. Mobile Genetic Element Flexibility as an Underlying Principle to Bacterial Evolution. Annu Rev Microbiol 77, 603–624 (2023).

3. Christie, G. E. & Dokland, T. Pirates of the Caudovirales. Virology 434, 210–221 (2012).

4. Lindqvist, B. H., Dehò, G. & Calendar, R. Mechanisms of genome propagation and helper exploitation by satellite phage P4. Microbiol Rev 57, 683–702 (1993).

5. Chen, J. & Novick, R. P. Phage-Mediated Intergeneric Transfer of Toxin Genes. Science 323, 139– 141 (2009).

6. Fillol-Salom, A. et al. Phage-inducible chromosomal islands are ubiquitous within the bacterial universe. ISME J 12, 2114–2128 (2018).

7. Barth, Z. K., Netter, Z., Angermeyer, A., Bhardwaj, P. & Seed, K. D. A Family of Viral Satellites Manipulates Invading Virus Gene Expression and Can Affect Cholera Toxin Mobilization. mSystems 5, e00358–20 (2020).

8. Ibarra-Chávez, R., Hansen, M. F., Pinilla-Redondo, R., Seed, K. D. & Trivedi, U. Phage satellites and their emerging applications in biotechnology. FEMS Microbiol Rev 45, fuab031 (2021).

9. Penadés, J. R., Chen, J., Quiles-Puchalt, N., Carpena, N. & Novick, R. P. Bacteriophage-mediated spread of bacterial virulence genes. Current Opinion in Microbiology 23, 171–178 (2015).

10. Eppley, J. M., Biller, S. J., Luo, E., Burger, A. & DeLong, E. F. Marine viral particles reveal an expansive repertoire of phage-parasitizing mobile elements. Proceedings of the National Academy of Sciences 119, e2212722119 (2022).

11. Moura de Sousa, J. A. & Rocha, E. P. C. To catch a hijacker: abundance, evolution and genetic diversity of P4-like bacteriophage satellites. Philos Trans R Soc Lond B Biol Sci 377, 20200475 (2022).

12. Penadés, J. R. & Christie, G. E. The Phage-Inducible Chromosomal Islands: A Family of Highly Evolved Molecular Parasites. Annual Review of Virology 2, 181–201 (2015).

13. de Sousa, J. A. M., Fillol-Salom, A., Penadés, J. R. & Rocha, E. P. C. Identification and characterization of thousands of bacteriophage satellites across bacteria. Nucleic Acids Res 51, 2759–2777 (2023).

14. Mai-Prochnow, A. et al. ‘Big things in small packages: the genetics of filamentous phage and effects on fitness of their host’. FEMS Microbiology Reviews 39, 465–487 (2015).

15. Hay, I. D. & Lithgow, T. Filamentous phages: masters of a microbial sharing economy. EMBO Rep 20, e47427 (2019).

16. Waldor, M. K. & Mekalanos, J. J. Lysogenic conversion by a filamentous phage encoding cholera toxin. Science 272, 1910–1914 (1996).

17. Webb, J. S. et al. Cell death in Pseudomonas aeruginosa biofilm development. J Bacteriol 185, 4585–4592 (2003).

18. Webb, J. S., Lau, M. & Kjelleberg, S. Bacteriophage and phenotypic variation in Pseudomonas aeruginosa biofilm development. J Bacteriol 186, 8066–8073 (2004).

19. Rice, S. A. et al. The biofilm life cycle and virulence of Pseudomonas aeruginosa are dependent on a filamentous prophage. ISME J 3, 271–282 (2009).

20. Addy, H. S., Askora, A., Kawasaki, T., Fujie, M. & Yamada, T. Utilization of Filamentous Phage ϕRSM3 to Control Bacterial Wilt Caused by Ralstonia solanacearum. Plant Dis 96, 1204–1209 (2012).

21. Kreienbaum, M. et al. Isolation and Characterization of Shewanella Phage Thanatos Infecting and Lysing Shewanella oneidensis and Promoting Nascent Biofilm Formation. Front Microbiol 11, 573260 (2020).

22. Wawrzyniak, P., Płucienniczak, G. & Bartosik, D. The Different Faces of Rolling-Circle Replication and Its Multifunctional Initiator Proteins. Frontiers in Microbiology 8, (2017).

23. Touchon, M., Moura de Sousa, J. A. & Rocha, E. P. Embracing the enemy: the diversification of microbial gene repertoires by phage-mediated horizontal gene transfer. Curr Opin Microbiol 38, 66–73 (2017).

24. Cleary, J. M. & Ray, D. S. Replication of the plasmid pBR322 under the control of a cloned replication origin from the single-stranded DNA phage M13. Proceedings of the National Academy of Sciences 77, 4638–4642 (1980).

25. Levinson, A., Silver, D. & Seed, B. Minimal size plasmids containing an M13 origin for production of single-strand transducing particles. J Mol Appl Genet 2, 507–517 (1984).

26. Peeters, B. P., Schoenmakers, J. G. & Konings, R. N. Plasmid pKUN9, a versatile vector for the selective packaging of both DNA strands into single-stranded DNA-containing phage-like particles. Gene 41, 39–46 (1986).

27. Dotto, G. P., Enea, V. & Zinderi, N. D. Functional analysis of bacteriophage f1 intergenic region. Virology 114, 463–473 (1981).

28. Dotto, G. P. & Zinder, N. D. The morphogenetic signal of bacteriophage f1. Virology 130, 252–256 (1983).

29. Ruiz-Masó, J. A. et al. Plasmid Rolling-Circle Replication. Microbiology Spectrum 3, 3.1.16 (2015).

30. Zhang, Y. et al. MazF cleaves cellular mRNAs specifically at ACA to block protein synthesis in Escherichia coli. Mol Cell 12, 913–923 (2003).

31. Culviner, P. H. & Laub, M. T. Global Analysis of the E. coli Toxin MazF Reveals Widespread Cleavage of mRNA and the Inhibition of rRNA Maturation and Ribosome Biogenesis. Mol Cell 70, 868–880.e10 (2018).

32. Harms, A., Brodersen, D. E., Mitarai, N. & Gerdes, K. Toxins, Targets, and Triggers: An Overview of Toxin-Antitoxin Biology. Mol Cell 70, 768–784 (2018).

33. Jurėnas, D., Fraikin, N., Goormaghtigh, F. & Van Melderen, L. Biology and evolution of bacterial toxin-antitoxin systems. Nat Rev Microbiol 20, 335–350 (2022).

34. LeRoux, M. & Laub, M. T. Toxin-Antitoxin Systems as Phage Defense Elements. Annu. Rev. Microbiol. 76, 21–43 (2022).

35. Lerner, T. J. & Model, P. The “steady state” of coliphage f1: DNA synthesis late in infection. Virology 115, 282–294 (1981).

36. Gescher, J. S., Cordova, C. D. & Spormann, A. M. Dissimilatory iron reduction in Escherichia coli: identification of CymA of Shewanella oneidensis and NapC of E. coli as ferric reductases. Mol Microbiol 68, 706–719 (2008).

37. Wick, R. R., Judd, L. M., Gorrie, C. L. & Holt, K. E. Unicycler: Resolving bacterial genome assemblies from short and long sequencing reads. PLOS Computational Biology 13, e1005595 (2017).

38. Seemann, T. Prokka: rapid prokaryotic genome annotation. Bioinformatics 30, 2068–2069 (2014).

39. Jumper, J. et al. Applying and improving AlphaFold at CASP14. Proteins 89, 1711–1721 (2021).

40. Terzian, P., et al. PHROG: families of prokaryotic virus proteins clustered using remote homology. NAR Genomics and Bioinformatics 3, lqab067 (2021).

41. Gibson, D. G. et al. Enzymatic assembly of DNA molecules up to several hundred kilobases. Nat Methods 6, 343–345 (2009).

42. Schmittgen, T. D. & Livak, K. J. Analyzing real-time PCR data by the comparative C(T) method. Nat Protoc 3, 1101–1108 (2008).

43. Bennett, N. J., Gagic, D., Sutherland-Smith, A. J. & Rakonjac, J. Characterization of a dual-function domain that mediates membrane insertion and excision of Ff filamentous bacteriophage. J Mol Biol 411, 972–985 (2011).

44. Beilstein, F. & Dreiseikelmann, B. Bacteriophages of freshwater Brevundimonas vesicularis isolates. Res Microbiol 157, 213–219 (2006).

45. Dreiseikelmann, B. et al. Characterization and genome comparisons of three Achromobacter phages of the family Siphoviridae. Arch Virol 162, 2191–2201 (2017).

46. Schindelin, J., et al. Fiji: an open-source platform for biological-image analysis. Nat Methods 9, 676–682 (2012).

