## Supplementary Tables S1-S4; Supplementary Figures 1-3 for "A rolling circle-replicating plasmid as an Inovirus phage satellite"

### – Supplementary information –

**Supplementary Table S1: Analysis of predicted Dolos and pDolos ORFs using Alphafold (Jumper et al, 2021), Foldseek (van Kempen et al, 2023), Protein BLAST (Altschul et al, 1997) and InterPro (Paysan-Lafosse et al, 2022).** Assignment of Dolos ORF function was mainly conducted by InterPro protein domain analysis, the amino acid (aa) length of proteins and the ORF order.

| ORF number | Alphafold structure* | [aa] | Foldseek analysis | Protein BLAST | InterPro | Assignment |
|---|---|---|---|---|---|---|
| **Phage Dolos** | | | | | | |
| **ORF 1** | 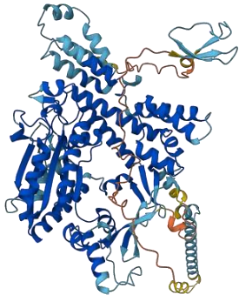 | 714 | - | Replication protein | | pII |
| **ORF 2** | 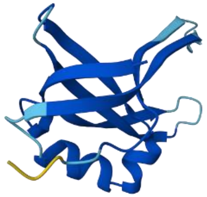 | 104 | - | | | pV |
| **ORF 3** | 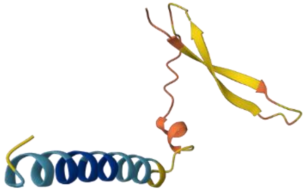 | 76 | - | Hypothetical protein | Transmembrane domain | pVII / pIX |

| | | | | | | |
|---|---|---|---|---|---|---|
| **ORF 4** | 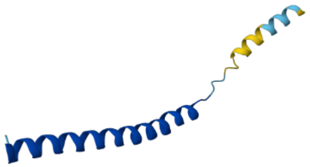 | 67 | - | Coat protein B | Transmembrane domain | pVIII |
| **ORF 5** | 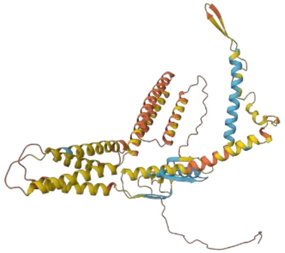 | 468 | - | Attachment protein (pIII) | Transmembrane domain, signalling domain | pIII |
| **ORF 6** | 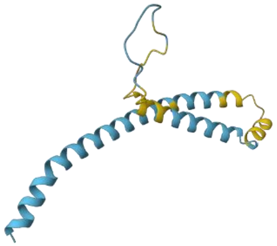 | 115 | - | gp6 | Transmembrane domain, signalling domain | pVI |
| **ORF 7** | 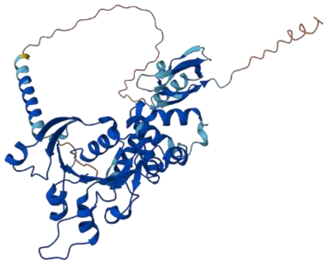 | 442 | - | Zot protein | Zot domain | pI |
| **ORF 8** | 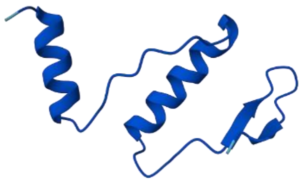 | 61 | Protein of unknown function | Hypothetical protein | no result | - |

| | | | | | | |
|---|---|---|---|---|---|---|
| **ORF 9** | 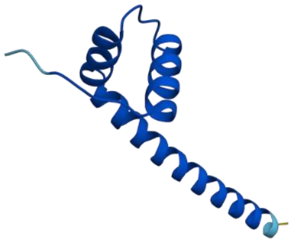 | 73 | Potential antitoxin (MazE family) [P0CL61] | NikA | no result | - |
| **ORF 10** | 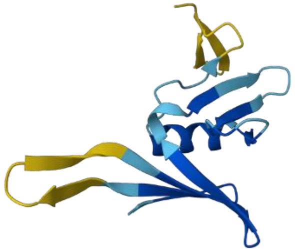 | 109 | Protein of unknown function | No result | Coil domain | - |
| **ORF 11** | 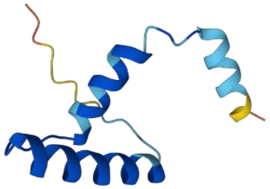 | 67 | Protein of unknown function | Hypothetical protein | No result | - |
| **pDolos** | | | | | | |
| **ORF 1** | 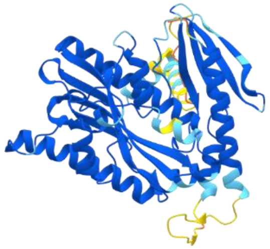 | 359 | Rep protein [P36229] | Rep protein | Rep domain | Rep protein |

| | | | | | | |
|---|---|---|---|---|---|---|
| **ORF 2** | 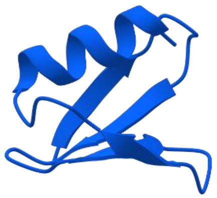 | 47 | Protein of unknown function | No result | No result | - |
| **ORF 3** | 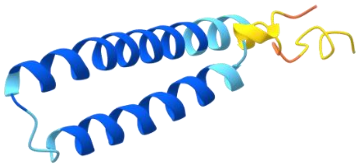 | 81 | Potential transcriptional repressor [X8FKK4] | No result | No result | - |
| **ORF 4** | 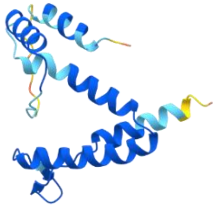 | 130 | Potential antitoxin (EndoAI family) [P96621] | No result | No result | - |
| **ORF 5** | 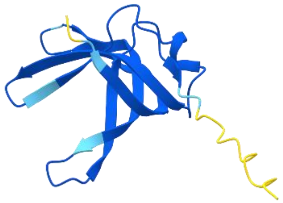 | 109 | Potential ssDNA binding protein [Q9PKZ4] | No result | No result | - |

*** model confidence:** dark blue: very high (pLDDT > 90), blue: confident (< 90 pLDDT > 70), yellow: low (< 70 pLDDT > 50), orange: very low (pLDDT < 50)

**Supplementary Table S2: List of bacterial species that can or cannot be infected by phage Dolos.** Infection was determined by spotting phage Dolos on agar overlay plates containing different bacterial species and subsequent spot-analysis after 48h of incubation.

| Not infected by Dolos | Infected by Dolos |
|---|---|
| *S. oneidensis* Isolate S17 | *S. oneidensis* MR-1 |
| *S. oneidensis* Isolate S52 | *S. oneidensis* MR-1 ΔλSo ΔMuSoII |
| *S. oneidensis* Isolate S54 | *S. baltica* Isolate S4 |
| *S. oneidensis* Isolate S62 | *S. baltica* Isolate S44 |
| *S. oneidensis* Isolate S63 | |
| *S. oneidensis* Isolate S66 | |
| *S. oneidensis* Isolate S69 | |
| *S. oneidensis* Isolate S74 | |
| *S. baltica* OS195 | |
| *S. baltica* Isolate S7 | |
| *S. baltica* Isolate S37 | |
| *S. baltica* Isolate S38 | |
| *S. baltica* Isolate S50 | |
| *S. amazonensis* SB2B | |
| *S. seohaensis* Isolate S8 | |
| *S. seohaensis* Isolate S10 | |
| *S. seohaensis* Isolate S11 | |
| *S. seohaensis* Isolate S31 | |
| *S. seohaensis* Isolate S32 | |
| *S. putrefaciens* CN-32 | |
| *S. putrefaciens* CN-32 ΔCas1_2 | |
| *S. putrefaciens* W3-18-1 | |
| *S. sp.* ANA-3 | |
| *S. sp.* MR-4 | |
| *S. sp.* MR-7 | |
| *E. coli* MG1655 | |
| *P. putida* KT2440 | |
| *V. cholerae* El Tor N16961 | |

**Supplementary Table 3: Bacterial strains and plasmids used in this study**

| Strain or plasmid | Description | Reference |
|---|---|---|
| ***Shewanella* strains** | | |
| *S. oneidensis* MR-1 | Wild type | Venkateswaran et al., 1999 |
| *S. oneidensis* MR-1 ΔLambdaSo ΔMuSo2 | Deletion of active prophages | Kreienbaum et al., 2020 |
| *S. oneidensis* Isolate S52 | | Jung-Schroers et al., 2017 |
| *S. oneidensis* Isolate S54 | | Jung-Schroers et al., 2017 |
| *S. oneidensis* Isolate S62 | | Jung-Schroers et al., 2017 |
| *S. oneidensis* Isolate S63 | | Jung-Schroers et al., 2017 |
| *S. oneidensis* Isolate S66 | | Jung-Schroers et al., 2017 |
| *S. oneidensis* Isolate S69 | | Jung-Schroers et al., 2017 |
| *S. oneidensis* Isolate S74 | | Jung-Schroers et al., 2017 |
| *S. baltica* Isolate S4 | | Jung-Schroers et al., 2017 |
| *S. baltica* Isolate S44 | | Jung-Schroers et al., 2017 |
| *S. baltica* Isolate S37 | | Jung-Schroers et al., 2017 |
| *S. baltica* Isolate S38 | | Jung-Schroers et al., 2017 |
| *S. baltica* Isolate S50 | | Jung-Schroers et al., 2017 |
| *S. amazonensis* SB2B | Wild type | Venkateswaran et al., 1998 |
| *S. seohaensis* Isolate S8 | | Jung-Schroers et al., 2017 |
| *S. seohaensis* Isolate S10 | | Jung-Schroers et al., 2017 |
| *S. seohaensis* Isolate S11 | | Jung-Schroers et al., 2017 |
| *S. seohaensis* Isolate S31 | | Jung-Schroers et al., 2017 |
| *S. seohaensis* Isolate S32 | | Jung-Schroers et al., 2017 |
| *S. putrefaciens* CN-32 | Wild type | Fredrickson et al., 1998 |
| *S. putrefaciens* CN-32 Δ*cas1_2* | Deletion of *cas1_2* | Dwarakanath et al. ,2015 |
| *S. putrefaciens* W3-18-1 | Wild type | Murray et al., 2001 |
| *S. sp.* ANA-3 | Wild type | Saltikov et al. 2003 |
| *S. sp.* MR-4 | Wild type | Nealson et al., 1991 |
| *S. sp.* MR-7 | Wild type | Nealson et al., 1991 |
| **Other strains** | | |
| *E. coli* MG1655 | K12 wild type | Jensen, 1993 |
| *P. putida KT2440* | Wild type | Nelson et al., 2002 |
| *V. cholerae* El Tor N16961 | Wild type | Heidelberg et al., 2000 |
| **Plasmids** | | |
| pDolos_K | pDolos harboring an additional kanamycin-resistance cassette ($Km^r$) | This work |
| pDolos_KV | pDolos_K also harboring *Venus* | This work |
| pDolos_KVL | pDolos_KV_also harboring *lacZ'* | This work |

**Supplementary Table 4: Primers used in this study**

| Identifier | Primer name | Sequence 5´→ 3´ |
|---|---|---|
| **pDolos constructs** | | |
| NES476 | MGE 179 cut FWD | CTAGAGGCTTGCAGAGCGA |
| NES477 | MGE 179 cut REV | AGGCCGCGCCCCGCCT |
| NES610 | MGE_Venus FWD | AATTCGAGCTCGGTACCCTTACTTGTACAGCTCGTCCATG |
| NES611 | MGE_Venus REV | ATCGAATTCCTGCAGCCCAGGAGGGCAAATATGGTGAGCA AGGGCGAG |
| NES612 | MGE_pBBR Promotor FWD | CTCGCCCTTGCTCACCATATTTGCCCTCCTGGGCTGCAGGA ATTCGAT |
| NES613 | MGE_pBBR Promotor REV | GCGATGGCCCACTACGTGTCATGCCGTTTGTGATGG |
| NES614 | MGE_Kan FWD | AAGCCATCACAAACGGCATGAGGAGCGCCTGAAGCCC |
| NES615 | MGE_Kan REV | GAATGATGTAGCCGTCAAGTTGTCATAACCTGAATCGCCA GCGG |
| NES616 | MGE_pBAD Promotor FWD | AATATGGTATTGATAATCCTTATGACAACTTGACGGCTAC |
| NES617 | MGE_pBAD Promotor REV | AGTGAATCCGTAATCATGGTCATATTTGCCCTCCTGGGTAC CGAGCTCGAATT |
| NES618 | MGE_LacZ FWD | CGAGCTCGGTACCCAGGAGGGCAAATATGACCATGATTAC GGATTCA |
| NES619 | MGE_LacZ REV | CGACTCTAGAGGATCCCCTTATTTTTGACACCAGACCAA |
| NES620 | MGE_Kan no_LacZ REV | GCCTGCAGGTCGACTCTAGAGGATCCCCGGATTATCAATA CCATATTTTTGAAA |
| NES486 | MGE_Kan Insert 179 FWD | TCAGGCGGGGCGCGGCCTCACGTAGTGGGCCATCG |
| NES487 | MGE_Kan Insert 179 REV | GCTCTGCAAGCCTCTAGGGATTATCAATACCATATTTTTGA AA |
| NES654 | Venus insertion MGE179 FWD | AGGCGGGGCGCGGCCTTTACTTGTACAGCTCGTCCATG |
| NES655 | LacZ insertion MGE179 REV | ATCGCTCTGCAAGCCTCTAGTTATTTTTGACACCAGACCAA CT |
| NES657 | no lacZ MGE179 REV | ATCGCTCTGCAAGCCTCTAGCCTGAATCGCCAGCGG |
| ***lacZ* check PCR** | | |
| NES625 | lacZ check 2 FWD | CCACCGATATTATTTGCCCG |
| NES619 | MGE_LacZ check REV | CGACTCTAGAGGATCCCCTTATTTTTGACACCAGACCAA |
| **qPCR** | | |
| NES581 | MGE qPCR FWD | CAGTTGGCCATTCATGGT |
| NES582 | MGE qPCR REV | GCAAGCAGATGATGAAACG |
| NES583 | MR-1 qPCR FWD | CTGTCTGAAACTCAACGGC |
| NES584 | MR-1 qPCR REV | CATGCCATTACTCTATCTCTACTGC |
| NES464 | Dolos qPCR FWD | TAGAACGATTCTCATGCTTCG |
| NES447 | Dolos qPCR REV | GGCTCGGTCTTTACTAAATGG |

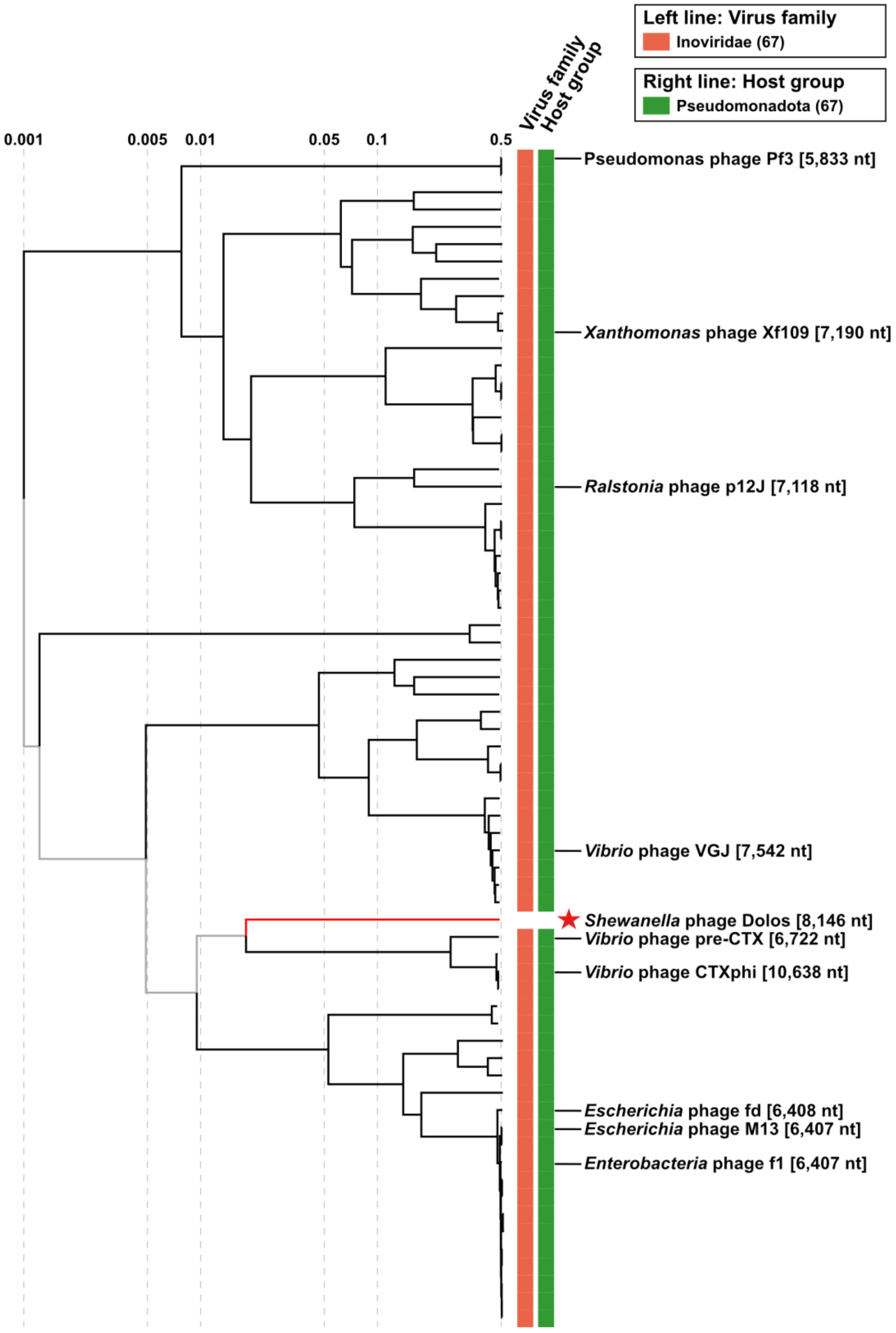

**Supplementary Figure 1: Viral proteomic tree.** The phylogenetic tree was generated using VipTree software (Nishimura et al, 2017). Phage Dolos and the genome sequences of all prokaryotic ssDNA viruses of the Virus-Host DB (RefSeq release 220) were included in the analysis. Left and right line are coloured according to the virus family or host group. Phage Dolos is highlighted with a red star.

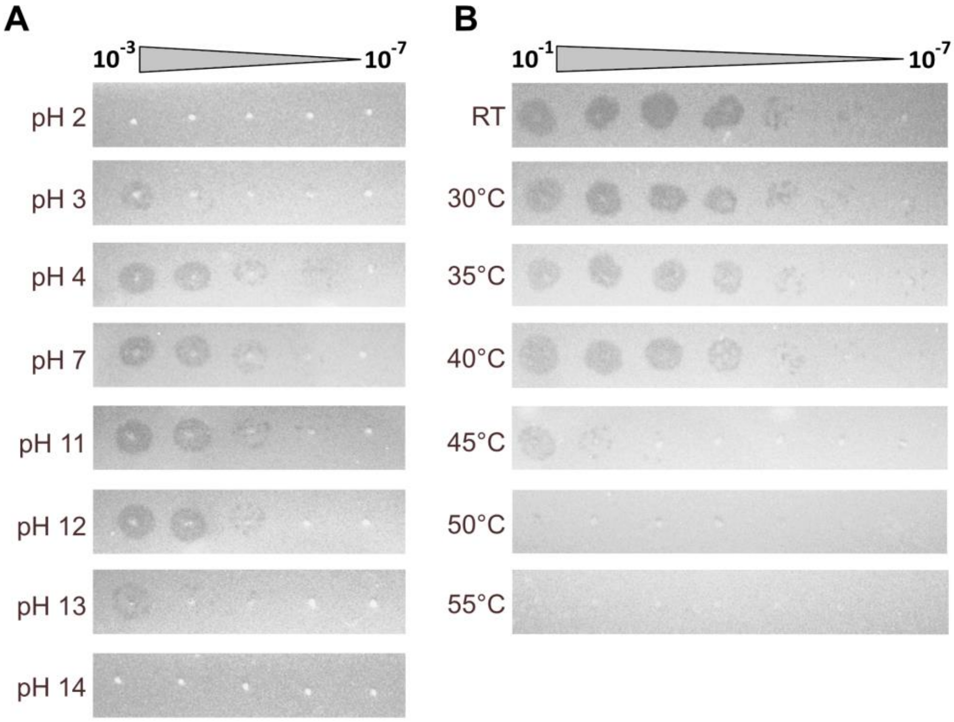

**Supplementary Figure 2: pH and temperature sensitivity of phage Dolos.** Serial dilutions of phage preparations exposed to different pH (**A**) or temperature (**B**) values (as indicated) for 24 h. The phage containing supernatant was then spotted on agar overlay plates using *S. oneidensis* ΔLambdaSo ΔMuSo2.

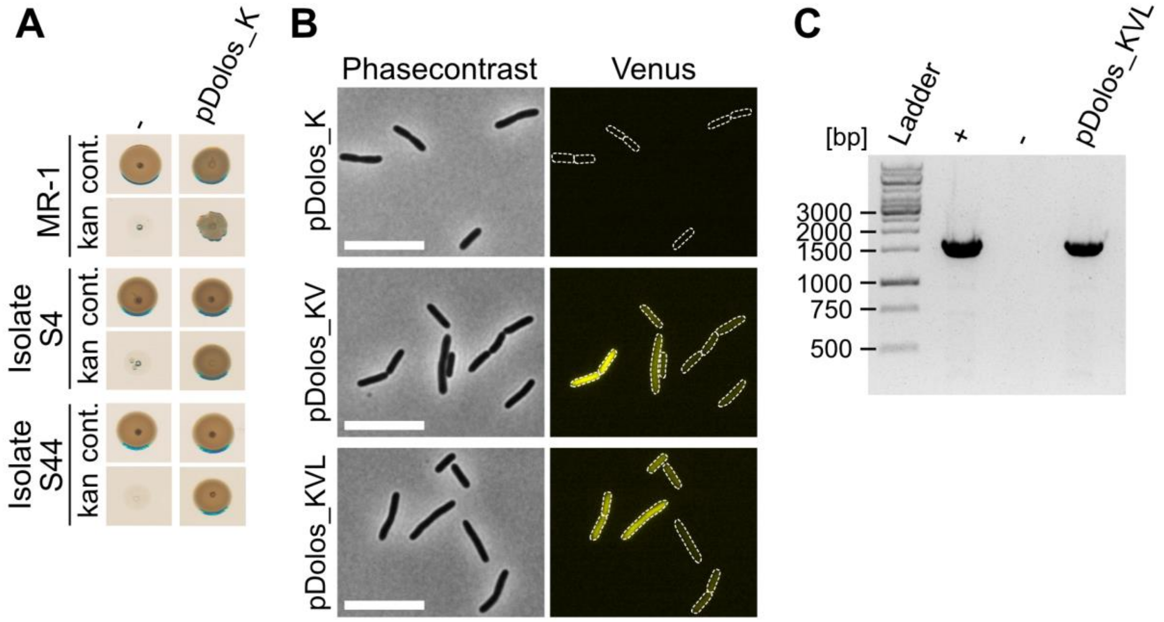

**Supplementary Figure 3: Lateral gene transfer of pDolos. A.** Transfer of pDolos_K into other *Shewanella* species. Bacteria were infected with an outgrown supernatant prepared from Dolos infected cells containing plasmid pDolos_K. 24h after transduction with the prepared infectious supernatant, cells ($OD_{600}$ 1) were spotted on 4M (cont.) and $4M_{Km}$ (kan) plates. Colony growth was imaged after 48h of incubation. **B and C.** Transduction of fluorophore Venus and gene *lacZ* by pDolos constructs. *S. oneidensis* ΔLambdaSo ΔMuSo2 cells were infected with an outgrown supernatant prepared from Dolos infected cells containing plasmid pDolos_K, pDolos_KV, or pDolos_KVL. 24h after infection mVenus fluorescence of cells ($OD_{600}$ 0.4) was analyzed via microscopy (**B**). Scale bar equals 5µm. Transmission of gene *lacZ* was analyzed by PCR followed by gel electrophoresis (**C**). -, *S. oneidensis* ΔLambdaSo ΔMuSo2 cells; +, isolated pDolos_KVL plasmid.

#### Additional References

Altschul SF, Madden TL, Schäffer AA, Zhang J, Zhang Z, Miller W, Lipman DJ. Gapped BLAST and PSI-BLAST: a new generation of protein database search programs. Nucleic Acids Res. 1997 Sep 1;25(17):3389-402. doi: 10.1093/nar/25.17.3389. PMID: 9254694; PMCID: PMC146917.

Dwarakanath S, Brenzinger S, Gleditzsch D, Plagens A, Klingl A, Thormann K, Randau L. Interference activity of a minimal Type I CRISPR-Cas system from *Shewanella putrefaciens*. Nucleic Acids Res. 2015 Oct 15;43(18):8913-23. doi: 10.1093/nar/gkv882. Epub 2015 Sep 8. PMID: 26350210; PMCID: PMC4605320.

Heidelberg JF, Eisen JA, Nelson WC, Clayton RA, Gwinn ML, Dodson RJ, et al. DNA sequence of both chromosomes of the cholera pathogen *Vibrio cholerae*. Nature. 2000 Aug 3;406(6795):477-83. doi: 10.1038/35020000. PMID: 10952301; PMCID: PMC8288016.

Jensen KF. The *Escherichia coli* K-12 "wild types" W3110 and MG1655 have an rph frameshift mutation that leads to pyrimidine starvation due to low *pyrE* expression levels. J Bacteriol. 1993 Jun;175(11):3401-7. doi: 10.1128/jb.175.11.3401-3407.1993. PMID: 8501045; PMCID: PMC204738.

Jumper J, Evans R, Pritzel A, Green T, Figurnov M, Ronneberger O, et al. Applying and improving AlphaFold at CASP14. Proteins. 2021 Dec;89(12):1711-1721. doi: 10.1002/prot.26257. PMID: 34599769; PMCID: PMC9299164.

Jung-Schroers V, Jung A, Ryll M, Bauer J, Teitge F, Steinhagen D. Methods for identification and differentiation of different *Shewanella* spp. isolates for diagnostic use. J Fish Dis. 2018 Apr;41(4):689-714. doi: 10.1111/jfd.12772. Epub 2017 Dec 27. PMID: 29280153.

Kreienbaum M, Dörrich AK, Brandt D, Schmid NE, Leonhard T, Hager F, et al. Isolation and Characterization of *Shewanella* Phage Thanatos Infecting and Lysing *Shewanella oneidensis* and Promoting Nascent Biofilm Formation. Front Microbiol. 2020 Sep 18;11:573260. doi: 10.3389/fmicb.2020.573260. PMID: 33072035; PMCID: PMC7530303.

Murray AE, Lies D, Li G, Nealson K, Zhou J, Tiedje JM. DNA/DNA hybridization to microarrays reveals gene-specific differences between closely related microbial genomes. Proc Natl Acad Sci U S A. 2001 Aug 14;98(17):9853-8. doi: 10.1073/pnas.171178898. Epub 2001 Aug 7. PMID: 11493693; PMCID: PMC55542.

Nealson, KH, Myers, CR, Wimpee, BB. Isolation and identification of manganese-reducing bacteria and estimates of microbial Mn(IV)-reducing potential in the Black Sea. Deep Sea Res Part A. Oceanograph Res Papers, 1991, 38, 907-920.

Nelson KE, Weinel C, Paulsen IT, Dodson RJ, Hilbert H, Martins dos Santos VA, et al. Complete genome sequence and comparative analysis of the metabolically versatile *Pseudomonas putida* KT2440. Environ Microbiol. 2002 Dec;4(12):799-808. doi: 10.1046/j.1462-2920.2002.00366.x. Erratum in: Environ Microbiol. 2003 Jul;5(7):630. PMID: 12534463.

Nishimura Y, Yoshida T, Kuronishi M, Uehara H, Ogata H, Goto S. ViPTree: the viral proteomic tree server. Bioinformatics. 2017 Aug 1;33(15):2379-2380. doi: 10.1093/bioinformatics/btx157. PMID: 28379287.

Paysan-Lafosse T, Blum M, Chuguransky S, Grego T, Pinto BL, Salazar GA, et al. InterPro in 2022. Nucleic Acids Res. 2023 Jan 6;51(D1):D418-D427. doi: 10.1093/nar/gkac993. PMID: 36350672; PMCID: PMC9825450.

Saltikov CW, Cifuentes A, Venkateswaran K, Newman DK. The ars detoxification system is advantageous but not required for As(V) respiration by the genetically tractable *Shewanella* species strain ANA-3. Appl Environ Microbiol. 2003 May;69(5):2800-9. doi: 10.1128/AEM.69.5.2800-2809.2003. PMID: 12732551; PMCID: PMC154534.

van Kempen M, Kim SS, Tumescheit C, Mirdita M, Lee J, Gilchrist CLM, Söding J, Steinegger M. Fast and accurate protein structure search with Foldseek. Nat Biotechnol. 2023 May 8. doi: 10.1038/s41587-023-01773-0. Epub ahead of print. PMID: 37156916.

Venkateswaran K, Dollhopf ME, Aller R, Stackebrandt E, Nealson KH. *Shewanella amazonensis* sp. nov., a novel metal-reducing facultative anaerobe from Amazonian shelf muds. Int J Syst Bacteriol. 1998 Jul;48 Pt 3:965-72. doi: 10.1099/00207713-48-3-965. PMID: 9734053.

Venkateswaran K, Moser DP, Dollhopf ME, Lies DP, Saffarini DA, MacGregor BJ, et al. Polyphasic taxonomy of the genus *Shewanella* and description of *Shewanella oneidensis* sp. nov. Int J Syst Bacteriol. 1999 Apr;49 Pt 2:705-24. doi: 10.1099/00207713-49-2-705. PMID: 10319494.
